# Systems-level analysis identifies key regulators driving epileptogenesis in temporal lobe epilepsy

**DOI:** 10.1101/688069

**Authors:** Yingxue Fu, Zihu Guo, Ziyin Wu, Liyang Chen, Yaohua Ma, Zhenzhong Wang, Wei Xiao, Yonghua Wang

## Abstract

Temporal lobe epilepsy (TLE) is the most prevalent and often devastating form of epilepsy. The molecular mechanism underlying the development of TLE remains largely unknown, which hinders the discovery of effective anti-epileptogenic drugs. In this study, we built a systems-level analytic framework which integrates gene meta-signatures, gene coexpression network and cellular regulatory network to unveil the evolution landscape of epileptogenic process and to identify key regulators that govern the transition between different epileptogenesis stages. The time-specific hippocampal transcriptomic profiles from five independent rodent TLE models were grouped into acute, latent and chronic stages of epileptogenesis, and were utilized for generating stage-specific gene expression signatures. 13 cell-type specific functional modules were identified from the epilepsy-context coexpression network, and five of them were significantly associated with the entire epileptogenic process. By inferring the differential protein activity of gene regulators in each stage, 265 key regulators underlying epileptogenesis were obtained. Among them, 122 regulators were demonstrated being associated with high seizure frequency and/or hippocampal sclerosis in human TLE patients. Importantly, we discovered four new gene regulators (*ANXA5, FAM107A, SEPT2* and *SPARC*) whose upregulation may drive the process of epileptogenesis and further lead to chronic recurrent seizures or hippocampal sclerosis. Our findings provide a landscape of the gene network dynamics underlying epileptogenesis and uncovered candidate regulators that may serve as potential targets for future anti-epileptogenic therapy development.

## Introduction

Epilepsy is a complex neurological disorder characterized by recurrent unprovoked seizures, of which temporal lobe epilepsy (TLE) is the most prevalent form [1]. The term epileptogenesis refers to the gradual process through which normal neuronal networks are altered resulting in the generation of chronic spontaneous seizures [2, 3]. The process can be triggered by diverse brain insults, including traumatic brain injury, stroke, infections and prolonged seizures such as status epilepticus (SE), and is typically thought to involve three stages [4, 5]. The first is the acute phase right after the brain insult, in which a cascade of morphologic and biologic changes occurs in the injured area. This is followed by a variable latent period during which behavioral seizures are not observed. The third stage is chronic, established epilepsy with the emergence of spontaneous seizures. Identifying the multiple dysregulated gene regulators that contribute to epileptogenesis in TLE is crucial for developing effective anti-epileptogenic drugs [6]. Several large-scale molecular signaling cascades such as mTOR, BDNF-TrkB and REST/NRSF pathways, have been demonstrated playing a role in epileptogenesis [7-9]. However, the detailed molecular mechanisms underlying the evolution process of epileptogenesis remain largely unknown.

The presence of various high-throughput omics technologies offers a great opportunity to unveil the molecular and cellular dynamics underlying epileptogenesis. Recently, a large-scale transcriptomic profiling of surgically resected hippocampi from TLE patients has been generated and used to identify gene-regulatory networks and regulators genetically associated with epilepsy [10, 11]. However, there are obvious limitations and challenges in exploring the process of epileptogenesis in human epileptic tissues. One drawback is that omics studies of human TLE generally lack appropriate control samples of healthy brain tissues. Furthermore, the specimens collected from hippocampus surgery for TLE patients are usually at an advanced stage and have been subjected to the treatment of various antiepileptic drugs (AEDs) [12]. Alternatively, well-characterized animal TLE models which mimic prominent histopathological and electroencephalographic features of human TLE can be employed to examine the key molecular alterations during epileptogenesis [13]. Only a few reports have studied the genome-wide molecular changes throughout epileptogenesis using animal TLE models [14, 15]. While other studies covered time points more closely related with either acute responses to SE or cumulative effects of chronic spontaneous seizures [16, 17]. As the modeling approaches and tissue dissection time varies across these studies, a systematic integration analysis of the existing datasets will likely provide a more comprehensive and robust molecular profiling for the epileptogenic process from the early hippocampal injury to the onset of chronic epilepsy.

Systems biology-based approaches that utilize network theory to organize transcriptome datasets have been used to prioritize candidate disease genes or to discern transcriptional regulatory programs [18-20]. One method to infer critical genes (hubs) and gene set–phenotype associations from gene expression data is the coexpression network analysis, which builds scale-free gene networks based on the pairwise gene expression correlations [21]. Genes with higher similarity scores tend to co-activate in a specific biological condition. Although coexpression analysis can help identify genes or gene modules that associate with the disease or biological phenotypes, it normally does not infer causality or distinguish between regulatory and regulated genes [22]. The algorithm for the reconstruction of accurate cellular networks (ARACNe) uses an information theoretic approach to eliminate most indirect interactions inferred by co-expression methods, leaving those expected to be regulatory [23]. Although originally being applied to infer the relationship between transcription factors and their target genes, the method can also be adapted to infer the indirect transcriptional targets for other kind of regulators, such as signaling proteins [24]. Using these methods, the observed gene expression changes can be placed into a systems context that was related to the underlying disease biology.

In this study, we proposed a systems-level analytic framework which integrates gene meta-signatures, gene coexpression network and cellular regulatory network to reveal the evolution landscape of epileptogenic process and distinguish key regulators that govern the transition between different epileptogenesis stages (**Fig. 1**). The time-specific hippocampal transcriptomic profiles of rodent TLE models were collected and classified into acute, latent and chronic stages of epileptogenesis. These profiles were then utilized for generating stage-specific gene expression signatures. Functional modules were detected from the coexpression network and their association with each epileptogenesis stage was assessed. Further, key gene regulators underlying epileptogenesis were identified by inferring the differential protein activity of regulators in each stage compared to control group. The influence of key regulators on synaptic signaling pathways were also explored. Finally, the validity of these key regulators was proved by their association with seizure frequency and hippocampal sclerosis in human TLE patients.

**Fig. 1.**
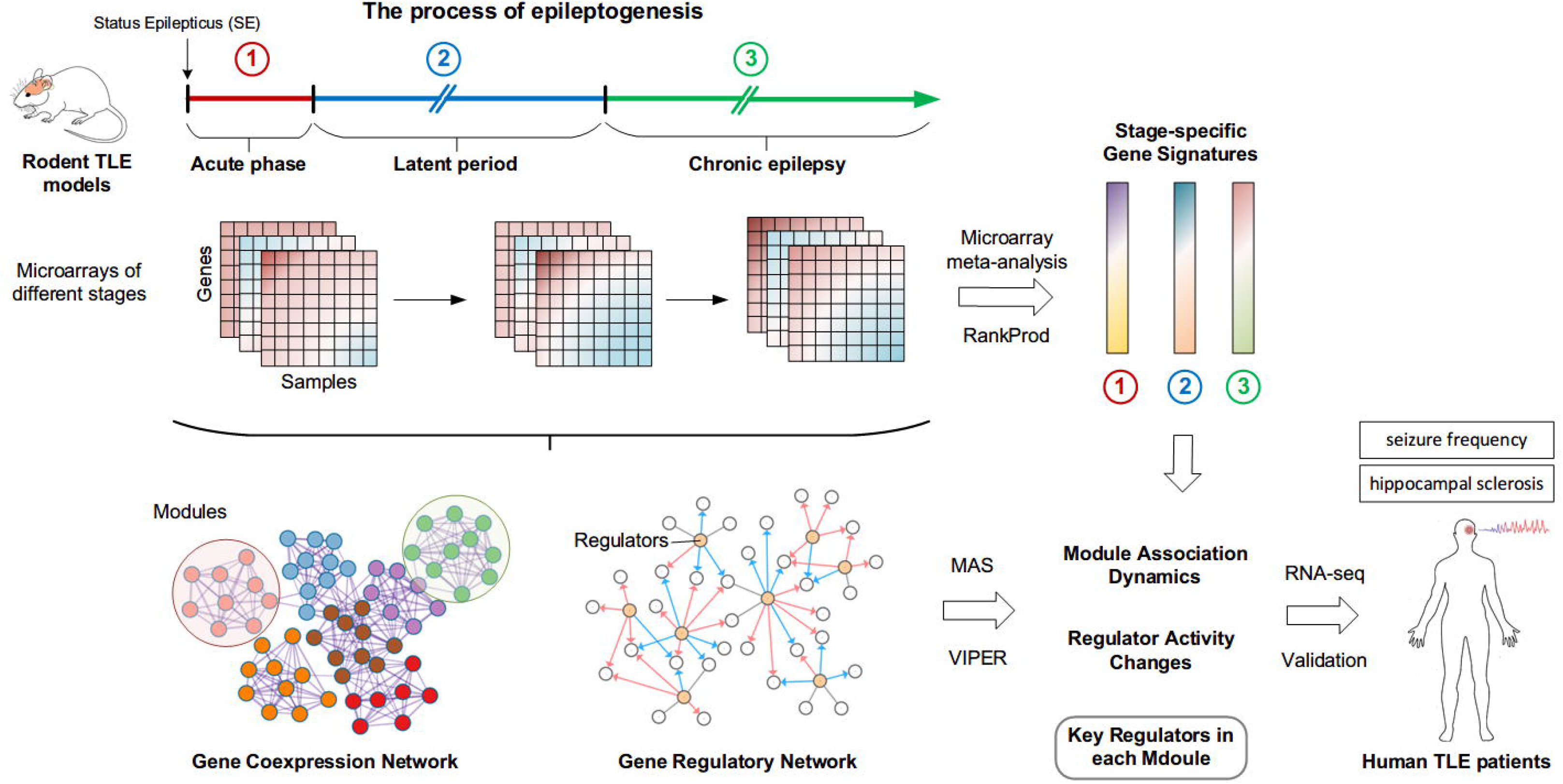
Schematic overview of the study design. To investigate the molecular mechanisms underlying the epileptogenic process, we proposed an analytic framework which comprises four steps. Firstly, the hippocampal transcriptome datasets of rodent TLE models were collected and divided into the acute, latent and chronic stages based on the described tissue extraction time and phenotypes. Meta-analysis was then performed using the RankProd method to evaluate differential gene expression and to generate the gene meta-signature for each stage. Secondly, to elucidate the functional organization of genes under the epilepsy context, gene coexpression network was constructed and used for identifying functional modules. A module association score (MAS) was then defined to quantify a module’s association degree with an epileptogenesis stage. Thirdly, to identify key regulators controlling the transition between stages, a gene regulatory network was constructed and the VIPER algorithm was employed to infer regulator activity changes in epileptic samples. Finally, using the RNA-seq profiles of human TLE patients with recurrent seizures and hippocampal sclerosis, key regulators associated with both conditions were screened out.

## Methods

### Data collection and preprocessing

We searched the Gene Expression Omnibus (GEO) database using key words “temporal lobe epilepsy”, “TLE” or “MTLE”, and restricted the study type as “Expression profiling by array, or by high throughput sequencing”. The organisms of samples were limited to Homo sapiens, Rattus norvegicus and Mus musculus. After manually checking all resulting datasets, we obtained five microarray datasets of rodent TLE models that covered different time points following SE and two human TLE patient RNA-seq datasets with epilepsy symptom information (seizure frequency or hippocampal sclerosis). Details about these datasets, including accession numbers, platforms and references, were listed in **Table 1**. For microarray datasets, the series matrix files were downloaded and then subjected to quality assessment using the *arrayQualityMetrics* package from Bioconductor [25]. Outliers were identified using heatmaps and dendrograms based on inter-array expression distances, and also boxplots and density estimate plots. For samples from GSE27166 and GSE73878, which contain both sides of hippocampus, only the expression profiles of the ipsilateral hippocampi were included. For RNA-seq datasets, the matrices of raw gene counts were downloaded from GEO database. Genes with very low counts across samples were filtered out based on the count-per-million (CPM) as implemented in the R package ‘edgeR’ (for detailed threshold, see the section “Human TLE patient RNA-seq data analysis”) [26]. To detect outlier across samples, the counts were normalized by the size factor of each library, and then log_2_ transformed and subjected to hierarchical clustering. Samples that did not show class-based clustering were removed in further analysis.

**Table 1.**
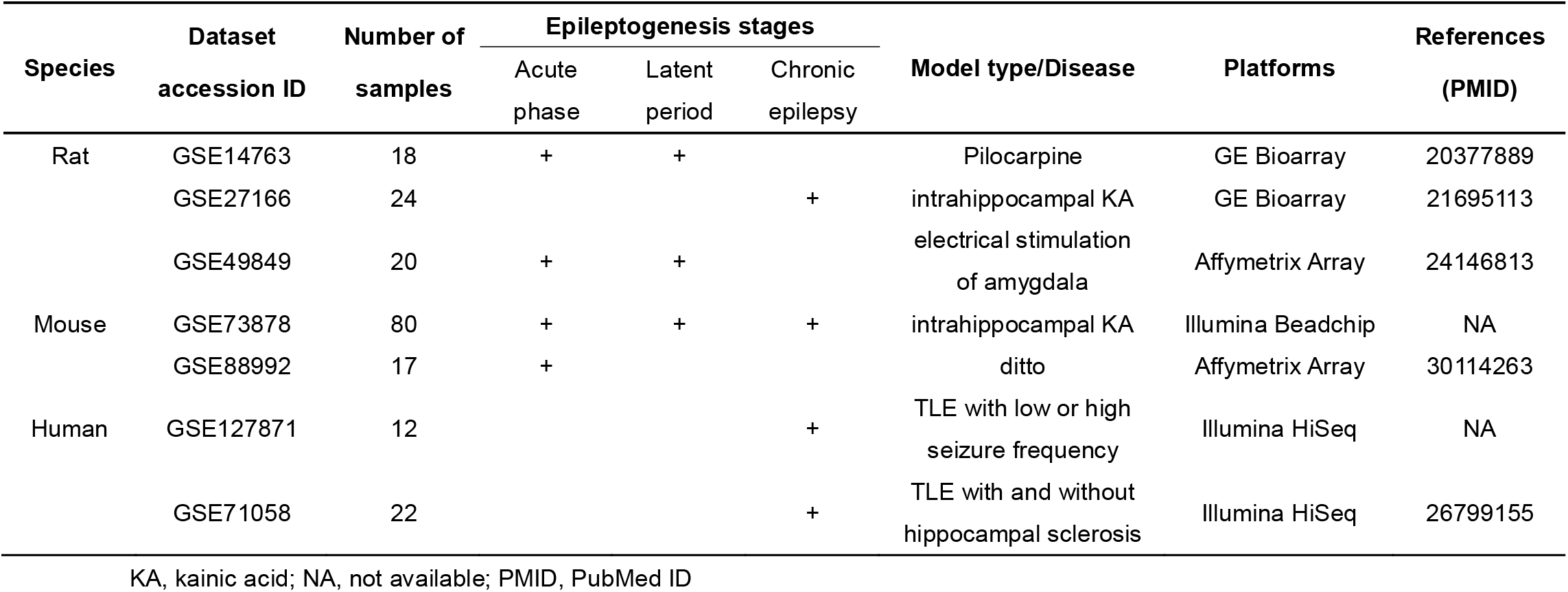
List of rodent TLE model microarray datasets used for epileptogenesis analysis and human TLE patient RNA-seq datasets used for validation.

### Principal component analysis

Unsupervised principal component analysis (PCA) was performed to further visualize the correlations among samples belonging to different epileptogenesis stages or epilepsy symptoms. All datasets were normalized and log2 transformed, and then analyzed using the *prcomp* function from the “stats” module in R. PCA methodology captures the inherent gene expression patterns in the data by projecting multivariate data objects onto a lower dimensional space while retaining much of the original variance [27].

### Differential gene expression analysis for individual datasets

For individual microarray datasets, we used the *limma* [28] and *RankProd* (RP) [29] packages from Bioconductor for differential expression analysis between sham control and epileptic samples of different epileptogenesis stages. The *limma* approach compare groups of samples by fitting gene-wise linear models and applying empirical Bayes methods to identify differentially expressed genes (DEGs). The genes with absolute log_2_FC (fold change)∟>0.5 and adjusted P-value for multiple comparison (FDR) <∟0.05 were considered significantly differentially expressed. RP is a non-parametric statistical method used to detect variables consistently upregulated or downregulated in replicate samples. It provides several advantages over linear modeling, including the biologically intuitive criterion, fewer model assumptions, and increased performance with noisy data. The DEGs were identified only based on the percentage of false predictions (pfp < 0.05) without any fold change restrictions. The list of DEGs identified by the two methods for each dataset was compared using Venn diagrams created by jvenn [30].

### Gene meta-signatures for specific epileptogenesis stages

For each epileptogenesis stage, the gene expression matrices of samples belonging to the corresponding stage were integrated from multiple datasets and meta-analysis was performed to evaluate the differential gene expression using the *RankProd* package [29]. Though the RP method was initially developed to detect DEGs in a single experiment, it is able to integrate datasets from multiple origins and overcome the heterogeneity among them because of the use of ranks instead of actual expression values. The four microarray platforms GPL1261, GPL2896, GPL6247 and GPL6885 for the rodent TLE model datasets contain 21720, 12733, 15124 and 17125 unique genes, respectively, of which 9139 genes were common across all platforms. The expression values of these common genes in each dataset were then extracted. For multiple probes that correspond to the same gene in a dataset, the probe with maximum mean expression values was retained to represent that gene. The RP method was then applied to the combined datasets of each epileptogenesis stage to assess the differential expression of genes. As RP employs separate ranks for up- and down-regulated genes, we integrated the two rank lists using the following equation (Eq. 1).

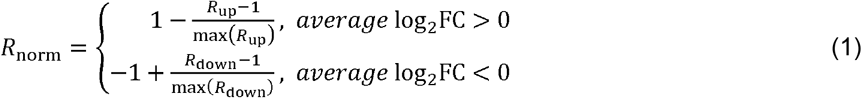

The gene lists with the normalized rank (*R*_norm_) and average log_2_FC was then served as gene meta-signatures for the three epileptogenesis stages.

### Gene coexpression network construction and module detection

To construct a epileptogenesis-context gene coexpression network, we subjected the dataset containing the entire process of epileptogenesis to weighted gene coexpression network analysis (WGCNA) [19, 21]. To overcome outlier bias, a robust correlation measure, biweight midcorrelation, was used to quantify the co-expression similarity *s_ij_* between each pair of genes [31]. Then, a weighted network adjacency matrix *A* = [*a_ij_*] was computed by applying a power function on all positive gene correlations, and was set to be zero when two genes have negative (or zero) correlations (Eq. 2).

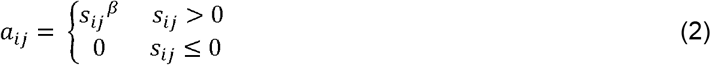

This ensures the connections of all gene pairs have the same direction and reduces the strength of weak correlations while preserving connection strength of highly correlated genes. The connectivity *k* for each gene was then defined as 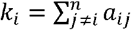 and was used for the network analysis. To balance the scale-free topology (i.e. *p*(*k*) ~ *k^−γ^*) and the sparsity of connections between genes in the network, a set of *β* values was evaluated to obtain the optimal specificity and sensitivity. To detect modules from the gene coexpression network, the topological overlap matrix (TOM), which reflects the relative interconnectedness between each pair of genes, was calculated. Based on the topological overlap dissimilarity (1–TOM) between genes, a gene dendrogram was generated using average hierarchical clustering. The dynamic branch cutting method was then used for detecting gene coexpression clusters (modules) in the dendrogram depending on its shape. Module eigengene, which is the first principal component of gene expression, was calculated to summarize the gene expression within a module and to merge modules with high similarities.

### Cell-type enrichment analysis

For the cell-type enrichment, marker genes of nine brain cell types were obtained from the single-cell RNA-seq profiles of the mouse cortex and hippocampus [32]. These include three types of neurons (cortical pyramidal neurons, CA1 pyramidal neurons and interneurons), four types of glia cells (astrocytes, oligodendrocytes, microglia and ependymal cells), and the vascular endothelial and mural cells. Enrichment between modules and the cell-type marker genes was measured using the hypergeometric test with subsequent BH correction for multiple comparison as implemented in the *userListEnrichment* function in the WGCNA package [21].

### Functional enrichment analysis

Functional meta-analysis for multiple gene sets (modules) was performed via Metascape [33] express analysis. Redundant terms were clustered into groups based on their similarities and the top 20 scored clusters were used as the final functional annotation for modules. Functional enrichment analysis against the KEGG pathway database for the key regulator list was performed using DAVID v6.8 [34].

### Module association score (MAS) with different epileptogenesis stages

To evaluate the association degree of modules with a specific epileptogenesis stage, a module association score (MAS) was defined to reflect the overrepresentation degree of a module at the extremes (top or bottom) of a ranked gene signature. For each module, only genes with the scaled intramodular connectivity greater than 0.2 were kept to represent the module. This reduced the noise for calculating MAS as genes with lower connectivity within a module typically contribute less to its functions. The MAS and the corresponding significance level were then calculated using the *fgsea* R package [35] against the gene meta-signature of each epileptogenesis stage. The *fgsea* method implements a special algorithm for fast gene set enrichment analysis. The significance of gene set enrichment was determined using the empirical enrichment score null distributions simultaneously calculated for all the gene set sizes. Module MAS with the adjusted P-value less than 0.05 was considered to be significantly associated with a specific epileptogenesis stage.

### Gene regulatory network construction

Candidate gene regulators were collected from three aspects. 1) Transcription factors or regulators (TFs). Transcription factors were obtained by extracting genes annotated in GO molecular function as GO:0003700, “DNA binding transcription factor activity”. Transcription regulators were the intersection of genes annotated as GO:0003677 “DNA binding” and genes annotated with GO:0140110, ‘transcription regulator activity’ or GO:0003612 “transcription coregulator activity” or GO:0006355, “regulation of transcription, DNA-templated”. 2) Synaptic proteins (SPs). SPs are regarded as genes annotated in GO cellular component as GO:0045202, ‘synapse’ or GO:0030424, “axon” or GO:0030425, “dendrite”. 3) Signaling proteins (Signal), which were built upon genes annotated in GO Biological Process GO:0007165 “signal transduction” and not overlapping with above two gene list. The genes corresponded to these GO terms were extracted using the *biomaRt* package [36]. These candidate regulators along with the gene expression matrix were subjected to the ARACNe-AP software [37] for reverse engineering a gene regulatory network. ARACNe was run with 100 bootstrap iterations with parameters set to 0 DPI (data processing inequality) tolerance and MI (mutual information) P-value threshold of 10^−8^.

### Protein activity inference for gene regulators

To infer the relative protein activity of the gene regulators at different epileptogenesis stages, we applied the VIPER algorithm [24] to test for regulon (a group of genes that are regulated by the same regulator) enrichment on stage-specific gene signatures. VIPER uses a probabilistic framework that integrates target mode of regulation (i.e., activated, repressed or undetermined represented by an index ranging from −1 to 1), statistical likelihoods of regulator-target interactions and target overlap between different regulators (pleiotropy). To compute the enrichment of a protein’s regulon in differentially expressed genes, an analytic rank-based enrichment analysis (aREA) method, which conduct a statistical analysis based on the mean of ranks, was used. The normalized enrichment score computed by aREA for each regulon were employed to quantitatively represent the relators’ relative protein activity in an epileptogenesis stage compared to the control group.

### Integration analysis of synaptic signaling pathways

To investigate the synaptic signaling variations at different epileptogenesis stages, an epilepsy-context synaptic signaling pathway was integrated and characterized. The integrated pathway is composed of key regulators enriched in the synapse-related pathways (i.e., the dopaminergic, cholinergic, glutamatergic, serotonergic and GABAergic synapses), and critical intracellular signaling pathways including MAPK signaling, calcium signaling, cGMP-PKG signaling and Ras signaling pathways. The interactions between these key regulators and their regulatory relationships with biological functions were extracted from the KEGG pathway database [38].

### Human TLE patient RNA-seq data analysis

For the dataset about TLE seizure frequency (GSE127871), genes with the CPM value greater than 0.4 in more than 50% of the samples were retained for further analysis. While for the hippocampal sclerosis dataset (GSE71058), the threshold was set to CPM > 0.1 in more than eight samples, which is the minimum number of samples in the two groups (with and without HS). After hierarchical clustering and PCA analysis for the samples in each dataset, the DESeq2 package [39] was used for differential expression analysis. Genes with the adjusted P-value less than 0.05 were considered as DEGs, and the log_2_FC ranked gene list were used as gene expression signatures. To make cross-species gene mappings, we standardized gene identifiers from microarray probe identifiers to NCBI Entrez ID identifiers and mapped mouse Entrez ID identifiers to their human ortholog using the *biomaRt* [36] package.

## Results

### Gene expression meta-signatures associated with different epileptogenesis stages

To investigate the molecular profiles underlying the epileptogenesis, we used the time-specific hippocampal transcriptome data of rodent TLE models from five independent studies (**Table 1**). These studies contain both rat and mouse TLE models and the modeling approaches were various, including systematic administration of pilocarpine, intrahippocampal KA injection and electrical stimulation of amygdala. The epileptic samples from these datasets covered a wide range of time points after SE, ranging from hours, to days and months. Based on the described tissue extraction time and phenotype, the samples in each dataset were divided into the control group and groups of three epileptogenesis stages, i.e., the acute phase, latent period and chronic epilepsy (**Fig. 1** and **Table 1**). After data normalization and preprocessing, 99 expression profiles were obtained and the individual datasets were then subjected to the PCA analysis. All datasets showed good separation among control samples and samples of different epileptogenesis stages along the first two PCs, of which PC1 accounted for the highest variation (27.8-49.8%) (**Fig. 2**).

**Fig. 2.**
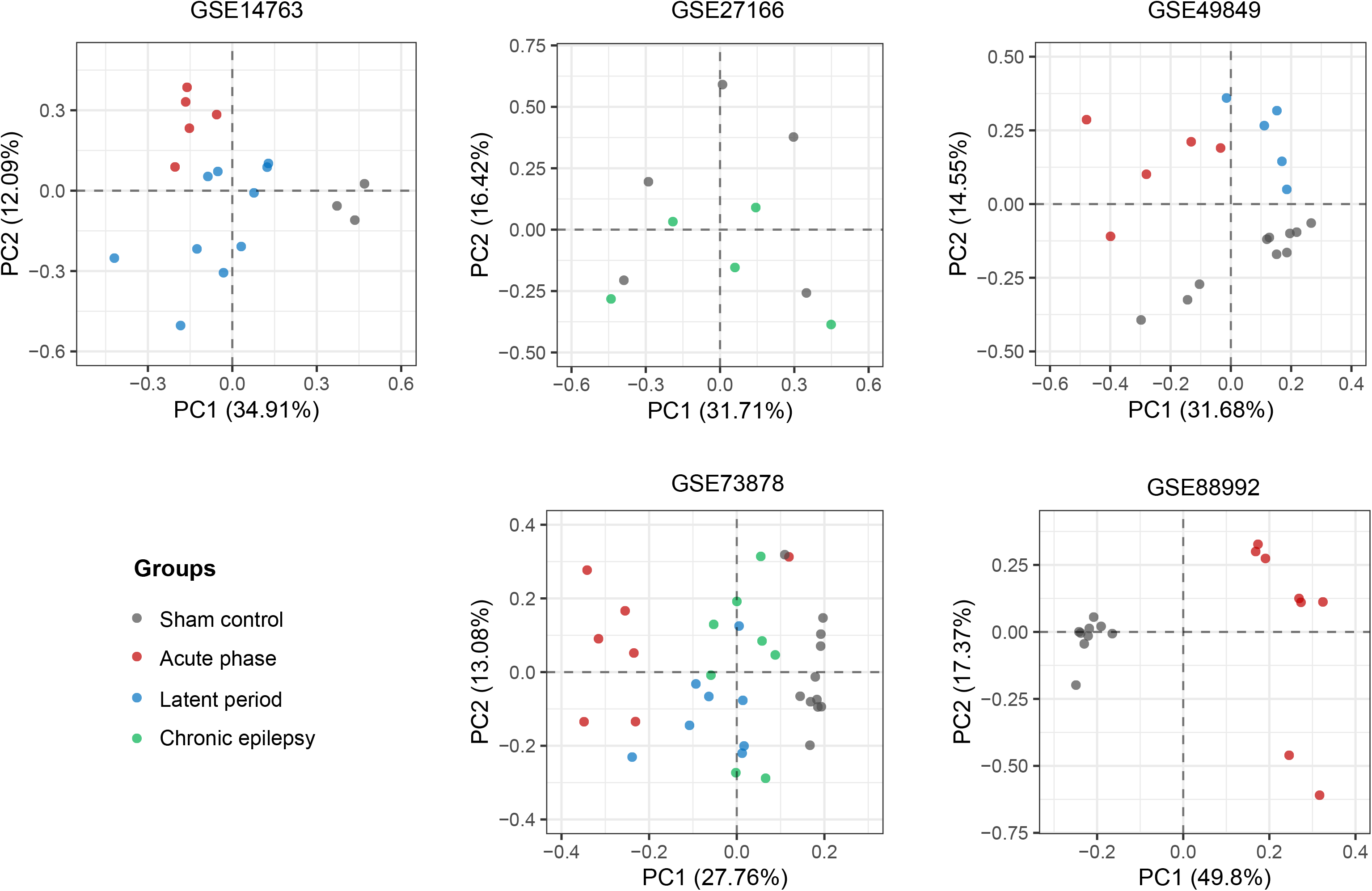
Principal component analysis (PCA) for microarrays of rodent TLE models. The variances captured by the first two PCs are shown along the respective axes. Samples belonging to the acute, latent and chronic stages are colored in red, blue and green, and control samples in gray. In all datasets, control samples and epileptic samples of different epileptogenesis stages form separate clusters.

To evaluate the differential gene expression between control samples and epileptic samples of each epileptogenesis stage, we first applied the *limma* method [28] to individual datasets. The differentially expressed genes (DEGs) were defined as those that achieved an absolute log_2_FC (fold change)∟>∟0.5 and a FDR∟<∟0.05 between control and epilepsy. There were four, three and two datasets that contain samples of the acute, latent and chronic stages, respectively. For the acute and latent stages, the differential expression analysis yielded gene lists with very small overlap across the datasets. While for the chronic phase, no DEGs were detected in one of the two datasets (**Supplementary Fig. S1a**). We then applied another differential expression analysis method RP [29] that differs from the linear modeling-based approaches. RP is a rank-based technique that detects genes that consistently appear among the most highly ranked genes (either strongly upregulated or downregulated) in a number of replicate samples. It identifies DEGs based on the estimated percentage of false predictions (pfp ∟ < ∟0.05). The RP-based method depicts a slightly better but still small overlap of DEGs among datasets (**Supplementary Fig. S1b**). These results suggest that direct comparison across individual datasets was not feasible due to the heterogeneity of experimental approaches and profiling platforms.

Since the RP algorithm transforms the actual expression values into ranks, it has the ability to handle variability among datasets and can be adapt to integrate datasets from multiple origins [40]. We thus adopted the RP for meta-analysis for each of the three epileptogenesis stages. We obtained a set of 2404, 1000 and 373 DEGs (pfp < 0.05) for the acute, latent and chronic stages, respectively. The top 100 DEGs of each stage were shown in **Supplementary Fig. S2**. It is evident from the heatmap that DEGs identified using the RP meta-analysis were consistently up- or down-regulated across most datasets. Among those top genes, 21 genes were dysregulated across all three epileptogenesis stages, including 14 upregulated genes (i.e. C3, *Cartpt, Cd44, Cd74, Cd9, Gfap, Ifitm3, Lcn2, Lgals3, Lyz2, Nptx2, Serping1, Timp1* and *Vim*) and 7 downregulated genes (i.e. *Bcl11a, Cdh8, Cygb, Fibcd1, Kctd4, Pkp2* and *Scn3b*), among which *Gfap* and *Scn3b* were widely described as biomarkers of the epileptogenesis [9, 41]. Besides, the mRNA expression of the “immediate early gene” (IEG) *Fos* and neural activity-dependent gene *Bdnf* was markedly induced in the acute and chronic stages, respectively, consistent with their reported roles in seizures and epilepsy [42]. Though the latent and chronic stages have less DEGs than the acute phase, this could be part of the result of different numbers of samples for each stage. As increased number of samples raises the power of the statistical test, leading to a higher number of selected genes. Therefore, we further extracted the whole RP-ranked gene list with genes’ average log_2_FC for each epileptogenesis stage, serving as the stage-specific gene expression signatures.

### Network functional dynamics during epileptogenesis

The gene meta-signatures associated with different epileptogenesis stages provide a significant starting point for dissecting the development of epilepsy. However, it is difficult to pinpoint epileptogenic mechanisms without considering the functional organization of these genes. To this aim, an epilepsy-context gene coexpression network was built using WGCNA [21] based on the dataset that contains samples of all three epileptogenesis stages. 13 gene modules (M1-M13), with their size varied from 44 to 1,896 genes, were identified (**Fig. 3a**). To investigate the cell-based context of these modules, we performed enrichment analysis against marker genes of nine brain cell types including neurons, glial cells and endothelial cells which were derived from single-cell RNA-seq analysis of the mouse cortex and hippocampus [32]. Module M1, the largest module with 1896 genes, was significantly enriched with three types of glial cells, among which microglia was the most enriched cell type (adjusted *P*-value = 4.0E-55) (**Supplementary Fig. S3**). Modules M3, 6, 7, 8 and 9 were specifically enriched for pyramidal neurons and/or interneurons. And modules M12 and 13 were enriched in the ependymal cell and mural cell.

**Fig. 3.**
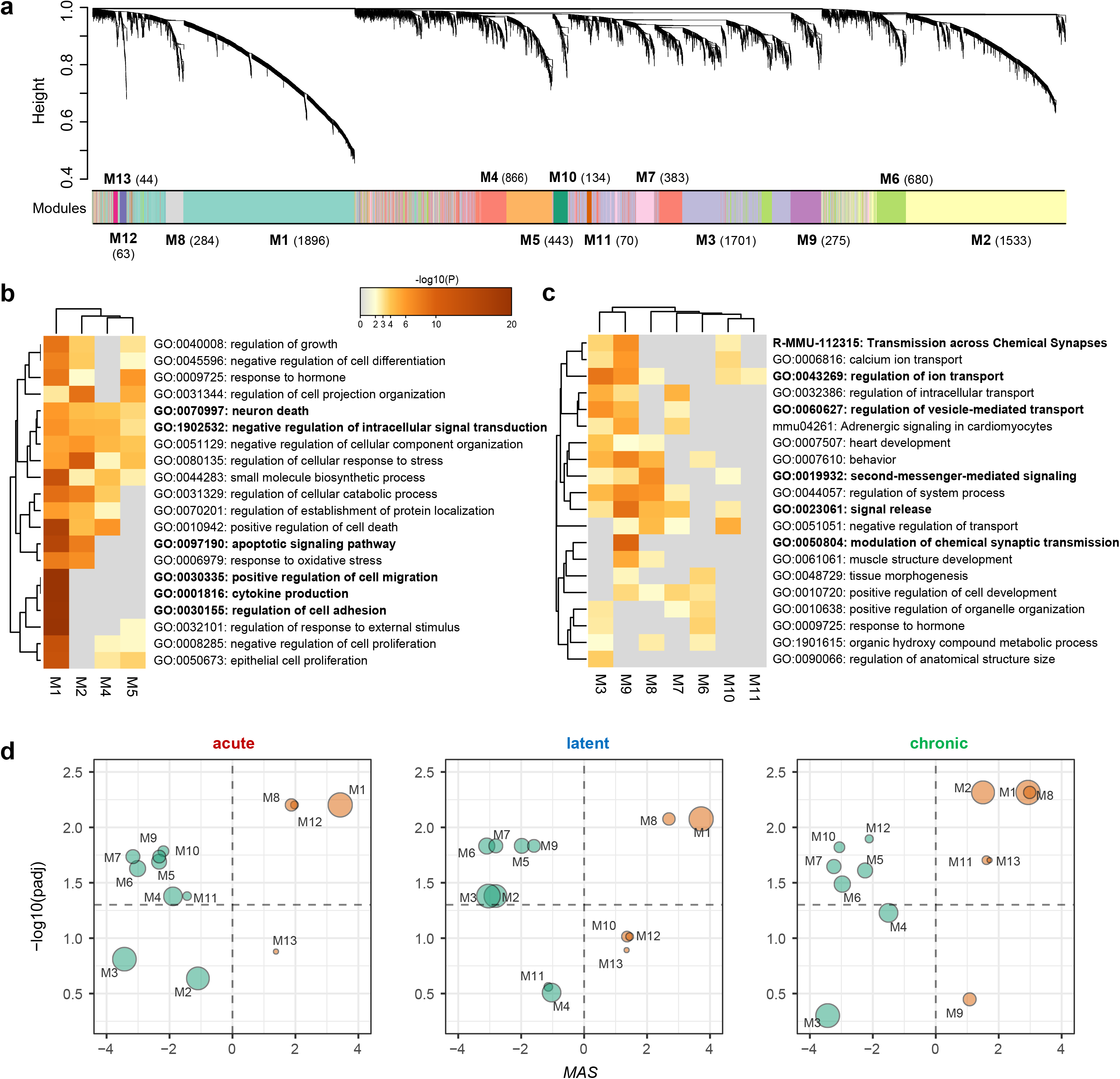
Network functional dynamics during epileptogenesis. **a.** Dendrogram showing clustering of 8,384 genes based on the topological overlap dissimilarity of genes in the gene coexpression network. Bottom color bar indicates the 13 gene coexpression modules (M1-M13) and their corresponding sizes (i.e. the number of genes in a module). **b** and **c.** Heatmaps showing the functional meta-analysis results for modules enriched with glia marker genes (M1, 2, 4 and 5) (**b**) and modules enriched with neuron marker genes (M3, 6, 7, 8, 9, 10 and 11) (**c**). The top 20 enriched functional terms are shown as rows and columns show modules. The heatmap is colored by the p-values. **d**. The MASs and corresponding significance levels (-log_10_(adjusted *P*-value)) of modules at the three epileptogenesis stages. The size of circle is proportional to the number of genes in each module used for calculating MAS.

Functional meta-analysis of these modules showed that module M1 mainly participated in GO biological processes of “positive regulation of cell migration”, “cytokine production”, “regulation of cell adhesion” and “apoptotic signaling pathway” (**Fig. 3b**). Moreover, M1 and modules M2, 4, 5 were all enriched with items of “neuron death” and “negative regulation of intracellular signal transduction”. Consistent with their enrichment for neuronal marker genes, modules M3, 6, 7, 8, 9, 10 and 11 were mainly involved in functions of the synapse, such as “regulation of ion transport”, “signal release”, “transmission across chemical synapses”, “regulation of vesicle-mediated transport” and “second-messenger-mediated signaling” (**Fig. 3c**).

To investigate how the expression of these modules was regulated at different epileptogenesis stages, we defined a module association score (MAS) to reflect the degree of which a module was enriched at the top or bottom of the stage-specific gene signatures. All modules were significantly associated with at least one epileptogenesis stage, among which five modules (M1, 8 and M5, 6, 7) were consistently up- or down-regulated in all three stages (**Fig. 3d**). The two positively associated modules M1 and 8 were mainly related to the inflammatory response and increased intracellular signaling activity. While the negatively associated modules M5, 6 and 7 may imply an impairment of the synapse function after SE. M4, 3 and 13 exhibited a specific association with the acute, latent and chronic stage, respectively. Module M9 was downregulated in the acute and latent phases but not the chronic epilepsy stage. Overall, these results provide a landscape of functional organization underlying epilepsy development, and also the functional dynamic changes during epileptogenesis.

### Identification of gene regulators driving epileptogenesis

To discover gene regulators controlling the transition from acute to latent and chronic stages of epileptogenesis, we interrogated the time-specific hippocampal transcriptome profiles using the VIPER algorithm [24] to infer the protein activity change of regulators in a specific stage. VIPER infers protein activity by systematically analyzing the expression of a protein’s regulon, which refers to the transcriptional targets of that protein. The ARACNe technique [37] which detect maximum information path targets was used to systematically infer regulons from epilepsy-specific gene expression data, resulting in an gene regulatory interactome of 41,364 interactions between 1,493 regulators and 5,695 target genes. VIPER then compute the enrichment of a protein’s regulon in differentially expressed genes based on a probabilistic framework that directly integrates target mode of regulation, regulator-target interaction confidence and target overlap between different regulators.

Differential protein activities of 521 regulators were obtained for all three epileptogenesis stages. For each stage, key regulators were defined as those with absolute differential protein activity score greater than two, which represents a significant activity alteration compared to the control group (**Fig. 4a**). There were 214, 198 and 156 key regulators respectively associated with the acute, latent and chronic stage (**Fig. 4b**). 43% of the key regulators were dysregulated in all three stages, indicating that these regulators were immediately involved in the epileptogenesis following SE, and exhibited continuing changes extending into the chronic epilepsy period. Modules M1, 3, 6, 7 and 8 have the highest numbers of key regulators (**Supplementary Fig. S4**). Regulator activities in both M1 and M8 were upregulated during the epileptogenesis, yet their activities exhibited opposite changing trends. Whereas M1 regulators were mainly associated with acute response after the SE, M8 regulators may play major roles in the latent and chronic epilepsy stages (**Fig. 4c**). Regulators in M3, 6 and 7 show constant downregulated activity at all three epileptogenesis stages.

**Fig. 4.**
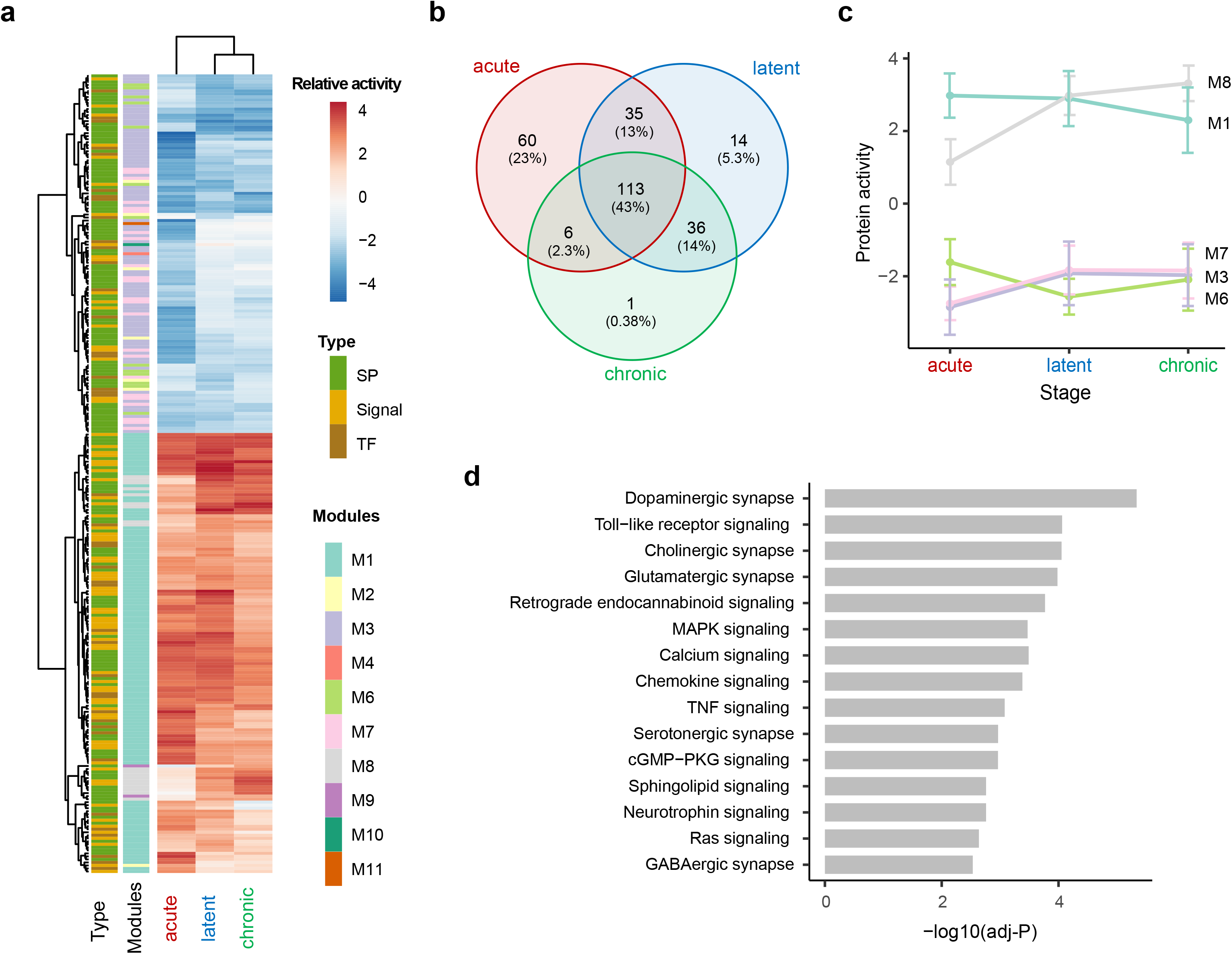
Identification of gene regulators driving epileptogenesis. **a.** Heatmap of the relative protein activity of 265 key regulators at the three epileptogenesis stages compared to the sham control group, with red denoting upregulation and blue denoting downregulation. The modules and types of these regulators are displayed at the left. SP, synaptic protein; Signal, signaling protein; TF, transcription factor. **b.** Venn plot showing the overlap of key regulators in the three epileptogenesis stages. **c.** Module-based regulator activity dynamics during epileptogenesis indicated by the mean and standard deviation of relative activities of key regulators in a module. **d.** KEGG pathway enrichment analysis for the 265 key regulators. The y-axis represents the top 15 significantly affected canonical pathways and x-axis the −log_10_ transformed BH adjusted P-values.

To understand the molecular processes affected by these key regulators, we performed enrichment analysis against the KEGG pathways database [38]. The analysis showed that the key regulators were involved in signaling pathways related to chemical synaptic transmission, immune response, growth factor signaling, and pathways related to cell proliferation and death (**Fig. 4d**). Among the synaptic transmission -related pathways, Dopaminergic synapse was the top enriched pathway (adjusted P-value∟=∟1.4E-05), followed by Cholinergic, Glutamatergic, Serotonergic and GABAergic synapses. Besides, the Retrograde endocannabinoid signaling, which can suppress both excitatory and some inhibitory synapses, was also enriched with the key regulators (adjusted P-value∟=∟1.7E-04). Endocannabinoids and their receptors are altered by epileptic seizures and can in turn control key epileptogenic circuits by inhibiting synaptic transmission in the hippocampus [43]. Multiple immune response-related pathways were also highly enriched, including the Toll-like receptor signaling, Chemokine signaling and TNF signaling pathways (adjusted P-value range∟=∟8.7E-05 to 8.3E-04). The Neurotrophin signaling pathway (adjusted P-value∟=∟1.7E-03) which can be activated by nerve growth factor (NGF) and brain-derived neurotrophic factor (BDNF) is an important pathway involved in the survival, development, and function of neurons. Other enriched pathways include the MAPK signaling, Calcium signaling, cGMP-PKG signaling and Ras signaling pathways (adjusted P-value range∟=∟3.4E-04 to 2.3E-03), which are critical intracellular signaling pathways related to multiple cellular functions.

### Variations of synaptic signaling at different epileptogenesis stages

A thorough knowledge of signaling pathways involved in both acute- and long-term responses to SE is crucial to unravel the origins of epilepsy [42]. To better understand the regulatory mechanism of synaptic transmission between neurons underlying epileptogenesis, we integrated and characterized an epilepsy-context synaptic signaling pathway that composed of the key regulators involved in synapse-related functions (**Fig. 5a**). A heatmap of the protein activities of these key regulators in different epileptogenesis stages was shown in **Fig. 5b**, in which the regulator types and modules were marked on the top.

**Fig. 5.**
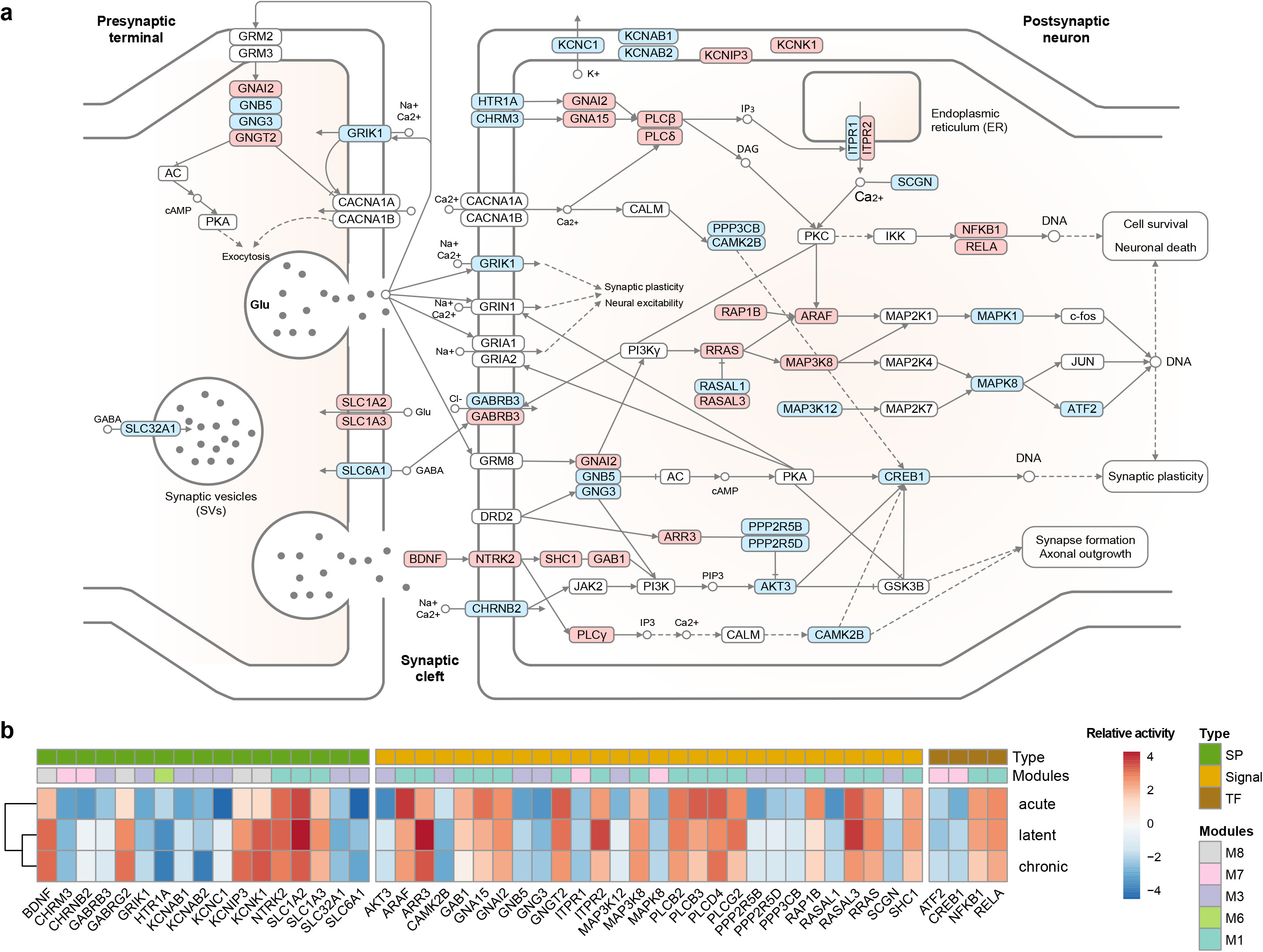
Variations of synaptic signaling at different epileptogenesis stages. **a.** The integrated synaptic signaling pathway showing the relationship between key regulators (KRs) and their activity changes during epileptogenesis (red, upregulated KR; blue, downregulated KR; white, other proteins). **b.** Heatmap showing differential protein activity of key regulators of the synaptic signaling pathway in three epileptogenesis stages. Dysregulated key regulators in the pathway are shown as rows and columns show differential protein activity in the acute, latent and chronic stage, respectively. The types and modules of these key regulators are shown on the top. SP, synaptic protein; Signal, signaling protein; TF, transcription factor.

We first examined the activity changes of key regulators located on synaptic membrane. For ionotropic glutamate receptors, only the kainate receptor (KAR) subunit 1 (*GRIK1*) was strikingly downregulated in the acute and latent stages. While regulators *GRIN1, GRIA1* and *GRIA2*, which are the subunits of NMDAR and AMPAR, also exhibited downregulation after SE, though not significant. Two subunits of GABA_A_ receptor (*GABRB3* and *GABRG2*) showed opposite changing trends during the epileptogenesis. *GABRB3* was significantly downregulated in the acute phase, while *GABRG2* was gradually upregulated in the latent and chronic periods. Besides, decreased activity of acetylcholine receptors (*CHRNB2* and *CHRM3*) and serotonin receptor (*HTR1A*) was also observed. Among these genes, mutations of *GRIN1, GABRB3, GABRG2* and *CHRNB2* have been reported associating with some familial epilepsy syndromes [44]. Notably, increased activity of glutamate transporters (*SLC1A2* and *SLC1A3*) and decreased activity of GABA transporters (*SLC6A1* and *SLC32A1*) further support the idea that imbalance between excitation and inhibition and altered threshold for neural excitation are underlying epileptic behaviors. The voltage-gated potassium channel Kv3.1 (*KCNC1*, alpha subunit) was markedly downregulated in the acute stage, implying the inability of the neuron to normally depolarize following SE. The accessory subunits of Kv1 (*KCNAB1* and *KCNAB2*) and Kv4 (*KCNIP3*) also display stage-specific activity changes. No significant variation was found for the activity of voltage-gated sodium or calcium channels.

Constant upregulation of *BDNF* and TrkB (*NTRK2*) was observed during the entire epileptogenesis, which further activated the adaptor proteins SHC1, GAB1 and PLCγ (*PLCG2*). This result is consistent with previous studies reporting that excessive activation of TrkB caused by SE promotes development of TLE [45, 46]. The two G_α_ subunits of G_i_ (*GNAI2*) and G_q_ (*GNA15*), which can inhibit adenylate cyclase (AC) and activate phospholipase C (PLC), exhibit significant upregulation, while the G_β_ and G_γ_ subunits (GNB5 and GNG3) of the G_βγ_ complex were downregulated strikingly in the acute stage. In accord with this, protein activity of PLC isotypes (*PLCB2, PLCB3*, and *PLCD4*) were also upregulated. This further activates the calcium signaling and protein kinase C (PKC), which can activate downstream transcription factors NFKB1 and RELA, inducing the transcription of target genes, like *c-fos* and *BCL-2*. For MAPK signaling, though there were increased activity of upstream Ras (RRAS), Raf (*ARAF*) and *MAP3K8*, both ERK (*MAPK1*) and JNK (*MAPK8*) activities were downregulated, along with the downregulation of activating transcription factor 2 (*ATF2*). Finally, the PI3K-AKT-CREB pathway, CaM kinase (CAMK2B) and calcium binding protein SCGN all exhibited decreased activity. In sum, these results demonstrate that the synapse-to-nucleus signaling underlying epileptogenesis were not simply up- or down-regulated, but displayed a complex restructured system related to neuronal hyperexcitation and impaired synaptic plasticity.

### Key regulators associated with seizure frequency and hippocampal sclerosis in human TLE

One of the direct outcomes of the epileptogenesis is the presence of spontaneous recurrent seizures. To investigate whether the identified key regulators were involved in controlling seizure frequency in patients with TLE, we utilized an RNA-Seq dataset of the hippocampal tissue resected from 12 medically intractable TLE patients with seizure frequencies ranging from 0.33 to 120 seizures per month. Hierarchical clustering and PCA analysis of the normalized profiles led to three clusters that can be regarded as the low (mean = 4.11 seizures/month), medium (mean = 13.2 seizures/month) and high (mean = 90 seizures/month) seizure frequency groups (**Supplementary Fig. 5a**). Differential expression analysis was then conducted between low or medium versus high SF groups to get both DEGs and gene expression signatures. Based on seizure frequency-associated gene signatures, we first tested whether the epileptogenesis-related functional modules were also disturbed in the higher seizure frequency group. The MAS calculated on low versus high SF gene expression signature demonstrated that module M1, which relate to acute inflammatory response, was the most strongly upregulated module, whereas modules M3, 5, 6, 7 and 9, which are related to synaptic transmission, were downregulated, in line with these modules’ expression changes in the epileptogenesis (**Fig. 6a**). For medium verse high SF gene expression changes, only modules M8 and M13 were significantly upregulated (**Fig. 6b**). As M8 and M13 were activated mainly in the chronic epilepsy stage, these results indicate that TLE patients with high seizure frequency indeed exhibit similar functional alterations detected on rodent TLE models undergo the epileptogenesis. We further examined whether the expression of key regulators was significantly altered in the high SF group, and found 108 (41%) key regulators overlap with DEGs detected in high versus low or medium SF groups.

**Fig. 6.**
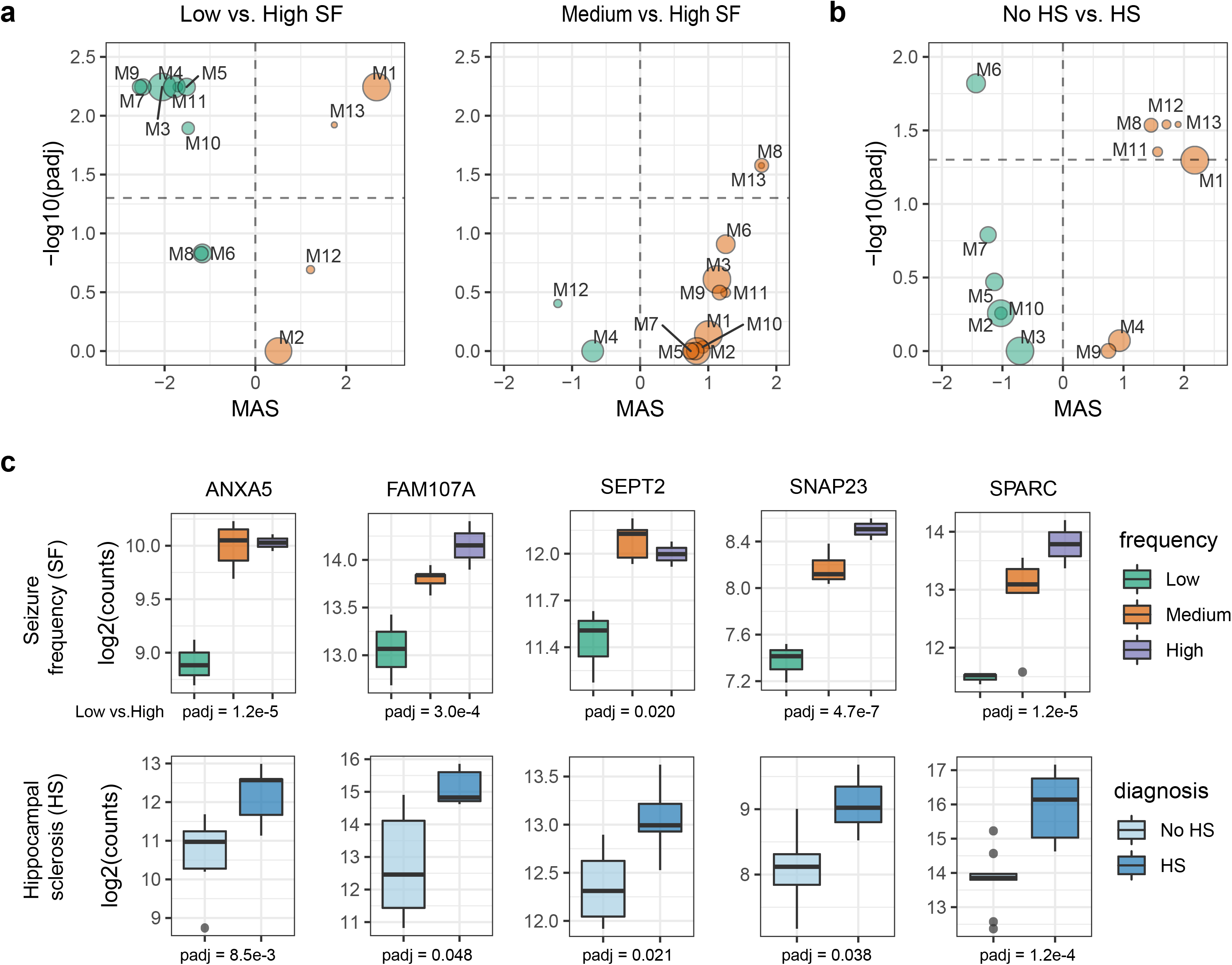
Modules and key regulators associated with seizure frequency and hippocampal sclerosis in human TLE. **a.** The MAS and corresponding significance level (-log_10_(adjusted *P*-value)) of modules calculated on gene expression signatures of low versus high seizure frequency (SF) group and medium versus high SF group. **b.** The same as **a** but calculated on the gene expression signature associated with hippocampal sclerosis (HS). In **a** and **b**, the size of circle is proportional to the number of genes in each module used for calculating the MAS. **c**. Boxplots showing the log_2_-scaled normalized counts of the five key regulators in TLE patients with different seizure frequency, and patients with and without HS. Bottom of each plot showing the adjusted P-values of genes derived from differential expression analysis using the DESeq2 package.

Hippocampal sclerosis (HS) is a common neuropathological condition encountered in TLE patients. It is featured with severe neuronal cell loss and gliosis in the hippocampus and can be both the cause and outcomes of the epileptogenesis [47]. Utilizing the RNA-seq profiles of dentate granule of MTLE patients with and without HS, we investigated how the epileptogenesis-associated functional modules and key regulators were modulated by HS (**Supplementary Fig. 5b**). Modules’ MASs on the HS-related gene signature showed that M1 had the highest MAS value (adjusted P-value = 0.0504). In addition, modules M8, 11, 12 and 13 were significantly upregulated and M6 was downregulated. 25 key regulators were found to be differentially expressed in patients with HS. By checking these regulators, we discovered 11 regulators that were associated with both hippocampal sclerosis and seizure frequency, namely ANXA5, ATF3, FAM107A, KCNK1, MAP7, NFIL3, RPS4X, SEPT2, SNAP23, SPARC and SV2B (**Fig. 6c** and **Supplementary Fig. S6**). Further analysis revealed that five of these key regulators (ANXA5, FAM107A, SEPT2, SNAP23 and SPARC) exhibit the same changing patterns (all upregulated) across the epileptogenesis and conditions of high seizure frequency and HS (**Fig. 6c**). Among them, SPARC (secreted protein acidic and rich in cysteine) was the most highly upregulated gene in both high SF (log_2_FC = 2.36, adj-P = 1.2E-05) and HS (log_2_FC = 2.24, adj-P = 1.2E-04) groups. Only SNAP23 (synaptosome associated protein 23), which is a vesical-associated protein, has been previously reported exhibiting upregulation in TLE patients with sclerotic hippocampus [48], while other four proteins have not been associated with epilepsy yet. Altogether, 122 out of 265 key regulators of epileptogenesis were associated with high seizure frequency and/or hippocampal sclerosis, indicating that our systems-level analysis has provided a valuable set of genes that may serve as potential therapeutic targets for epilepsy.

## Discussion

Epilepsy is a heterogeneous disorder with multiple origins and many different mechanisms of pathogenesis among patients [49]. To better understand the molecular mechanisms underlying epileptogenesis, it is necessary to take advantage of multiple animal epilepsy models with different origins and phenotypes. In this study, we performed an integration analysis on time-specific transcriptome profiles of various rodent TLE models, which include both chemical and electrical kindling models of epilepsy. Direct comparison of DEGs identified from individual datasets led to very limited information, indicating there need more systematic analysis to detect genes that are consistently up- or down-regulated across datasets of different origins. By applying a rank-based meta-analysis method RP to the gene expression matrix of each epileptogenesis stage, we obtained stage-specific gene expression signatures to depict the molecular features of the three epileptogenesis stages at a genome-wide level. Compared to using DEGs based on arbitrary cutoffs as gene signatures for a specific phenotype, gene lists with the relative fold change or rank information for each gene provide a more comprehensive molecular representation for each epileptogenesis stage.

Based on the stage-specific gene meta-signatures of epileptogenesis, we proposed a MAS value to assess the differences of cellular and molecular functions between stages. For example, the microglia-associated module M1, which is involved in multiple inflammation and immune response processes, was constantly upregulated throughout the entire epileptogenic process. This provides evidence to support the idea that inflammatory processes within the brain constitute a common and crucial mechanism in the pathophysiology of seizures and epilepsy [50]. Besides, we also observed that pyramidal neuron-enriched modules, M3 and M6, and interneuron-enriched module M7 were downregulated in all three epileptogenesis stages, which indicates that the synaptic transmission is severely impaired between these two types of neurons, leading to the excitation/inhabitation imbalance and circuit-level dysfunction in the hippocampus [3]. The consistency between the cell-type specificity and functional annotation of modules also demonstrates the biological significance of the identified modules in the context of epileptogenesis. The dynamic changes of modules’ association with different stages thus provide us a global landscape of the evolution process of epileptogenesis.

The expression dynamics of the functional modules are typically drove by defects in multiple gene regulators which exhibit concurrent and aberrant activities. For identifying key gene regulators, we inferred the differential protein activity of regulators in the three epileptogenesis stages. The regulator types include not only TFs, which are commonly regarded as the direct regulators controlling transition between different biological conditions [51], but also proteins on the synaptic membrane and intracellular signaling proteins. Given that various proteins located at the membrane of pre- or post-synapse are under the most directly impact when seizure activity occurs, the inclusion of synaptic and signaling proteins can help better depict the abnormalities of the synapse-to-nuclear signaling underlying epileptogenesis. Furthermore, the changing pattern of relative regulator activity in a module offers a straightforward illustration of how the regulators were modulated along with the development of epilepsy (**Fig. 4c**). For instance, we found that regulator activity of M8 exhibit gradual upregulation during epileptogenesis, implying these key regulators were remarkedly activated in the latent and chronic phases and may thus contribute to the formation of a brain state that supports recurrent, unprovoked seizures.

We further utilized transcriptome datasets of human TLE patients with or without hippocampal sclerosis and patients with different seizure frequencies to test the validity of key regulators detected from rodent TLE models. Hippocampal sclerosis is the most frequent cause of drug-resistant TLE, and presents a broad spectrum of electroclinical, structural and molecular pathology patterns [52]. We discovered four new gene regulators (ANXA5, FAM107A, SEPT2 and SPARC) from module M1 whose upregulation may contribute to higher seizure frequency and hippocampal sclerosis in TLE. Though the precise functions of these regulators are not fully understood, most of them are involved in the functions of the synapse and may have a potential role in maintaining synaptic plasticity. Detailed mechanisms of how these regulators drive the process of epileptogenesis and further lead to chronic recurrent seizures or hippocampal sclerosis need to be investigated using appropriate animal models of epilepsy in the future. In summary, our work provides a landscape of the gene network dynamics underlying epileptogenesis and highlighted candidate regulators controlling epileptogenesis that may warrant further investigation as potential anti-epileptogenic targets.

## Competing interests

The authors declare that they have no competing interests.

## Funding

This work was supported by the by the National Natural Science Foundation of China (NO. U1603285 and NO. 81803960)

## Authors’ contributions

Conceptualization: YXF; investigation: YXF; technical support: ZHG, ZYW, LYC and YHM; writing (original draft): YXF; writing (review and editing): YXF, ZHG, ZYW and YHW; supervision: ZZW and YHW; funding acquisition: YHW and WX. All authors read and approved the final manuscript.

## References

1. Moshe SL, Perucca E, Ryvlin P, Tomson T: Epilepsy: new advances. Lancet 2015, 385:884–898.

2. Pitkanen A, Lukasiuk K: Mechanisms of epileptogenesis and potential treatment targets. Lancet Neurol 2011, 10:173–186.

3. Goldberg EM, Coulter DA: Mechanisms of epileptogenesis: a convergence on neural circuit dysfunction. Nat Rev Neurosci 2013, 14:337–349.

4. Herman ST: Epilepsy after brain insult: targeting epileptogenesis. Neurology 2002, 59:S21–26.

5. Maguire J: Epileptogenesis: More Than Just the Latent Period. Epilepsy Curr 2016, 16:31–33.

6. Sloviter RS, Bumanglag AV: Defining “epileptogenesis” and identifying “antiepileptogenic targets” in animal models of acquired temporal lobe epilepsy is not as simple as it might seem. Neuropharmacology 2013, 69:3–15.

7. Citraro R, Leo A, Constanti A, Russo E, De Sarro G: mTOR pathway inhibition as a new therapeutic strategy in epilepsy and epileptogenesis. Pharmacol Res 2016, 107:333–343.

8. Heinrich C, Lahteinen S, Suzuki F, Anne-Marie L, Huber S, Haussler U, Haas C, Larmet Y, Castren E, Depaulis A: Increase in BDNF-mediated TrkB signaling promotes epileptogenesis in a mouse model of mesial temporal lobe epilepsy. Neurobiol Dis 2011, 42:35–47.

9. McClelland S, Brennan GP, Dube C, Rajpara S, Iyer S, Richichi C, Bernard C, Baram TZ: The transcription factor NRSF contributes to epileptogenesis by selective repression of a subset of target genes. Elife 2014, 3:e01267.

10. Johnson MR, Behmoaras J, Bottolo L, Krishnan ML, Pernhorst K, Santoscoy PLM, Rossetti T, Speed D, Srivastava PK, Chadeau-Hyam M, et al: Systems genetics identifies Sestrin 3 as a regulator of a proconvulsant gene network in human epileptic hippocampus. Nat Commun 2015, 6:6031.

11. Srivastava PK, van Eyll J, Godard P, Mazzuferi M, Delahaye-Duriez A, Steenwinckel JV, Gressens P, Danis B, Vandenplas C, Foerch P, et al: A systems-level framework for drug discovery identifies Csf1R as an anti-epileptic drug target. Nat Commun 2018, 9:3561.

12. Thom M: Review: Hippocampal sclerosis in epilepsy: a neuropathology review. Neuropathol Appl Neurobiol 2014, 40:520–543.

13. Mazzuferi M, Kumar G, Rospo C, Kaminski RM: Rapid epileptogenesis in the mouse pilocarpine model: video-EEG, pharmacokinetic and histopathological characterization. Exp Neurol 2012, 238:156–167.

14. Okamoto OK, Janjoppi L, Bonone FM, Pansani AP, da Silva AV, Scorza FA, Cavalheiro EA: Whole transcriptome analysis of the hippocampus: toward a molecular portrait of epileptogenesis. BMC Genomics 2010, 11:230.

15. Bot AM, Debski KJ, Lukasiuk K: Alterations in miRNA levels in the dentate gyrus in epileptic rats. PLoS One 2013, 8:e76051.

16. Kalozoumi G, Kel-Margoulis O, Vafiadaki E, Greenberg D, Bernard H, Soreq H, Depaulis A, Sanoudou D: Glial responses during epileptogenesis in Mus musculus point to potential therapeutic targets. PLoS One 2018, 13:e0201742.

17. Winden KD, Karsten SL, Bragin A, Kudo LC, Gehman L, Ruidera J, Geschwind DH, Engel J, Jr.: A systems level, functional genomics analysis of chronic epilepsy. PLoS One 2011, 6:e20763.

18. Ravasz E, Somera AL, Mongru DA, Oltvai ZN, Barabasi AL: Hierarchical organization of modularity in metabolic networks. Science 2002, 297:1551–1555.

19. Zhang B, Horvath S: A general framework for weighted gene co-expression network analysis. Stat Appl Genet Mol Biol 2005, 4:Article17.

20. Basso K, Margolin AA, Stolovitzky G, Klein U, Dalla-Favera R, Califano A: Reverse engineering of regulatory networks in human B cells. Nat Genet 2005, 37:382–390.

21. Langfelder P, Horvath S: WGCNA: an R package for weighted correlation network analysis. BMC Bioinformatics 2008, 9:559.

22. van Dam S, Vosa U, van der Graaf A, Franke L, de Magalhaes JP: Gene co-expression analysis for functional classification and gene-disease predictions. Brief Bioinform 2018, 19:575–592.

23. Margolin AA, Nemenman I, Basso K, Wiggins C, Stolovitzky G, Dalla Favera R, Califano A: ARACNE: an algorithm for the reconstruction of gene regulatory networks in a mammalian cellular context. BMC Bioinformatics 2006, 7 Suppl 1:S7.

24. Alvarez MJ, Shen Y, Giorgi FM, Lachmann A, Ding BB, Ye BH, Califano A: Functional characterization of somatic mutations in cancer using network-based inference of protein activity. Nat Genet 2016, 48:838–847.

25. Kauffmann A, Gentleman R, Huber W: arrayQualityMetrics--a bioconductor package for quality assessment of microarray data. Bioinformatics 2009, 25:415–416.

26. Robinson MD, McCarthy DJ, Smyth GK: edgeR: a Bioconductor package for differential expression analysis of digital gene expression data. Bioinformatics 2010, 26:139–140.

27. Yeung KY, Ruzzo WL: Principal component analysis for clustering gene expression data. Bioinformatics 2001, 17:763–774.

28. Ritchie ME, Phipson B, Wu D, Hu Y, Law CW, Shi W, Smyth GK: limma powers differential expression analyses for RNA-sequencing and microarray studies. Nucleic Acids Res 2015, 43:e47.

29. Del Carratore F, Jankevics A, Eisinga R, Heskes T, Hong F, Breitling R: RankProd 2.0: a refactored bioconductor package for detecting differentially expressed features in molecular profiling datasets. Bioinformatics 2017, 33:2774–2775.

30. Bardou P, Mariette J, Escudie F, Djemiel C, Klopp C: jvenn: an interactive Venn diagram viewer. BMC Bioinformatics 2014, 15:293.

31. Hardin J, Mitani A, Hicks L, VanKoten B: A robust measure of correlation between two genes on a microarray. BMC Bioinformatics 2007, 8:220.

32. Zeisel A, Munoz-Manchado AB, Codeluppi S, Lonnerberg P, La Manno G, Jureus A, Marques S, Munguba H, He L, Betsholtz C, et al: Brain structure. Cell types in the mouse cortex and hippocampus revealed by single-cell RNA-seq. Science 2015, 347:1138–1142.

33. Zhou Y, Zhou B, Pache L, Chang M, Khodabakhshi AH, Tanaseichuk O, Benner C, Chanda SK: Metascape provides a biologist-oriented resource for the analysis of systems-level datasets. Nat Commun 2019, 10:1523.

34. Huang da W, Sherman BT, Lempicki RA: Systematic and integrative analysis of large gene lists using DAVID bioinformatics resources. Nat Protoc 2009, 4:44–57.

35. Sergushichev A: An algorithm for fast preranked gene set enrichment analysis using cumulative statistic calculation. BioRxiv 2016:060012.

36. Durinck S, Spellman PT, Birney E, Huber W: Mapping identifiers for the integration of genomic datasets with the R/Bioconductor package biomaRt. Nat Protoc 2009, 4:1184–1191.

37. Lachmann A, Giorgi FM, Lopez G, Califano A: ARACNe-AP: gene network reverse engineering through adaptive partitioning inference of mutual information. Bioinformatics 2016, 32:2233–2235.

38. Kanehisa M, Goto S, Sato Y, Furumichi M, Tanabe M: KEGG for integration and interpretation of large-scale molecular data sets. Nucleic Acids Res 2012, 40:D109–114.

39. Love MI, Huber W, Anders S: Moderated estimation of fold change and dispersion for RNA-seq data with DESeq2. Genome Biol 2014, 15:550.

40. Hong F, Breitling R, McEntee CW, Wittner BS, Nemhauser JL, Chory J: RankProd: a bioconductor package for detecting differentially expressed genes in meta-analysis. Bioinformatics 2006, 22:2825–2827.

41. David Y, Cacheaux LP, Ivens S, Lapilover E, Heinemann U, Kaufer D, Friedman A: Astrocytic dysfunction in epileptogenesis: consequence of altered potassium and glutamate homeostasis? J Neurosci 2009, 29:10588–10599.

42. Bozzi Y, Dunleavy M, Henshall DC: Cell signaling underlying epileptic behavior. Front Behav Neurosci 2011, 5:45.

43. Monory K, Massa F, Egertova M, Eder M, Blaudzun H, Westenbroek R, Kelsch W, Jacob W, Marsch R, Ekker M, et al: The endocannabinoid system controls key epileptogenic circuits in the hippocampus. Neuron 2006, 51:455–466.

44. Fukata Y, Fukata M: Epilepsy and synaptic proteins. Curr Opin Neurobiol 2017, 45:1–8.

45. Gu B, Huang YZ, He XP, Joshi RB, Jang W, McNamara JO: A Peptide Uncoupling BDNF Receptor TrkB from Phospholipase Cgamma1 Prevents Epilepsy Induced by Status Epilepticus. Neuron 2015, 88:484–491.

46. Liu G, Gu B, He XP, Joshi RB, Wackerle HD, Rodriguiz RM, Wetsel WC, McNamara JO: Transient inhibition of TrkB kinase after status epilepticus prevents development of temporal lobe epilepsy. Neuron 2013, 79:31–38.

47. Walker MC: Hippocampal Sclerosis: Causes and Prevention. Semin Neurol 2015, 35:193–200.

48. Lee TS, Mane S, Eid T, Zhao H, Lin A, Guan Z, Kim JH, Schweitzer J, King-Stevens D, Weber P, et al: Gene expression in temporal lobe epilepsy is consistent with increased release of glutamate by astrocytes. Mol Med 2007, 13:1–13.

49. Watanabe S, Yamamori S, Otsuka S, Saito M, Suzuki E, Kataoka M, Miyaoka H, Takahashi M: Epileptogenesis and epileptic maturation in phosphorylation site-specific SNAP-25 mutant mice. Epilepsy Res 2015, 115:30–44.

50. Vezzani A, French J, Bartfai T, Baram TZ: The role of inflammation in epilepsy. Nat Rev Neurol 2011, 7:31–40.

51. Lefebvre C, Rajbhandari P, Alvarez MJ, Bandaru P, Lim WK, Sato M, Wang K, Sumazin P, Kustagi M, Bisikirska BC, et al: A human B-cell interactome identifies MYB and FOXM1 as master regulators of proliferation in germinal centers. Mol Syst Biol 2010, 6:377.

52. Blumcke I, Thom M, Aronica E, Armstrong DD, Bartolomei F, Bernasconi A, Bernasconi N, Bien CG, Cendes F, Coras R, et al: International consensus classification of hippocampal sclerosis in temporal lobe epilepsy: a Task Force report from the ILAE Commission on Diagnostic Methods. Epilepsia 2013, 54:1315–1329.

