## Supplementary Figures for "Systems-level analysis identifies key regulators driving epileptogenesis in temporal lobe epilepsy"

**Supplementary Information**

**Supplementary Fig. S1.** Venn diagrams showing the overlap of differentially expressed genes (DEGs) obtained from individual microarray datasets for each of the three epileptogenesis stages. **a.** The results from *limma* analysis, and **b.** The results from the individual *RankProd* (RP) analysis.

**Supplementary Fig. S2.** Heatmaps of the relative fold change of top 100 DEGs identified from RP meta-analysis for the three epileptogenesis stages. Datasets are shown as *columns* and genes are shown as *rows*. DEGs that are shared by all three epileptogenesis stages are marked in bold italic. Relative fold changes are shown with a pseudocolor scale (–1 to 1), with *red* denoting upregulation and *blue* denoting downregulation.

**Supplementary Fig. S3.** Cell-type enrichment analysis for modules. Nine brain cell types are shown in the rows. These include neurons (cortical pyramidal neurons, CA1 pyramidal neurons and interneurons), glia cells (astrocytes, oligodendrocytes, microglia and ependymal cells), and the vascular endothelial and mural cells. Modules were shown in the columns. Enrichment degree was determined using the hypergeometric test with subsequent BH correction for multiple testing (at FDR ≤ 0.05).

**Supplementary Fig. S4.** The distribution of the 265 key regulators (KRs) in the modules. Modules M1, 3, 6, 7 and 8 have the highest numbers of KRs, while modules M5, 12 and 13 do not have any KR. SP, synaptic protein; Signal, signaling protein; TF, transcription factor.

**Supplementary Fig. S5. a.** Hierarchical clustering of the two RNA-seq datasets of human TLE patients. Left, dataset for seizure frequency (GSE127871); Right, dataset for hippocampal sclerosis (GSE71058). Outliers that did not show class-based clustering were marked as red. **b.** PCA analysis for the remaining samples to further visualize the correlations among samples belonging to different groups.

**Supplementary Fig. S6.** Boxplots showing the log_2_-scaled normalized counts of the six key regulators, which exhibit opposite changing trends in TLE patients with different seizure frequency, and patients with and without HS. Bottom of each plot showing the adjusted P-values of genes derived from differential expression analysis using the DESeq2 package.
