## Supplementary figures and images for "Systems-level analysis identifies key regulators driving epileptogenesis in temporal lobe epilepsy"

### Supplementary Fig. S1

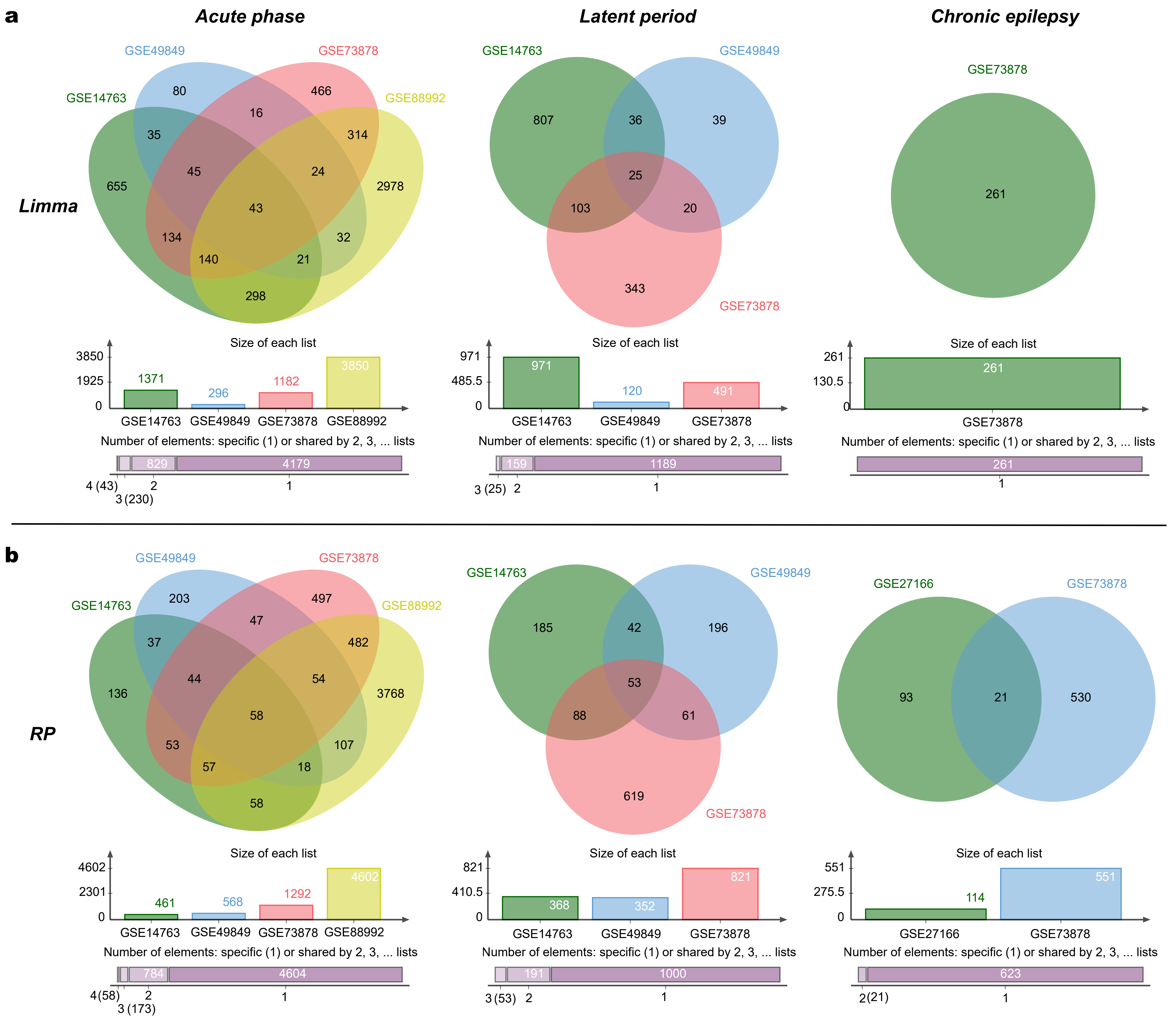

### Supplementary Fig. S2

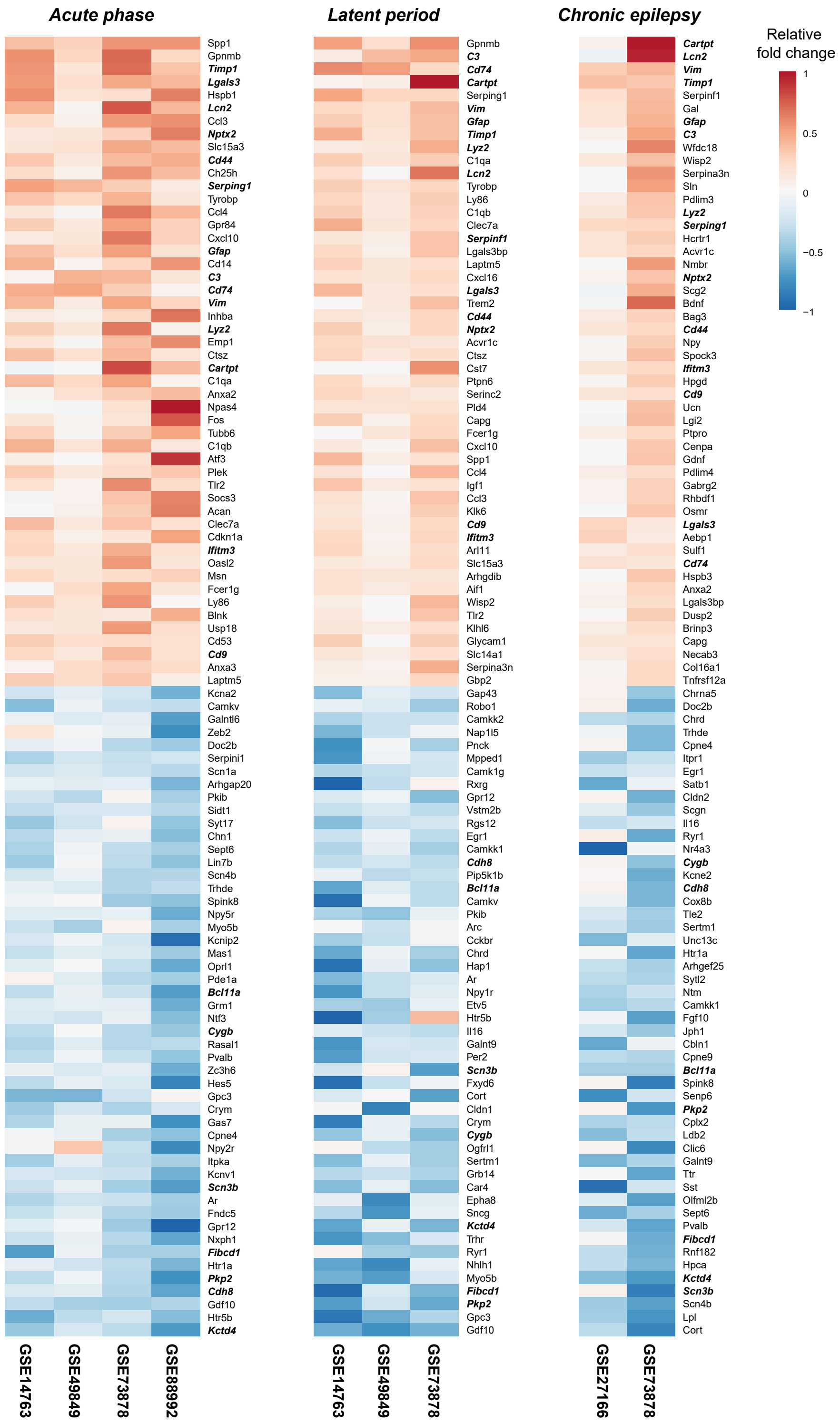

### Supplementary Fig. S3

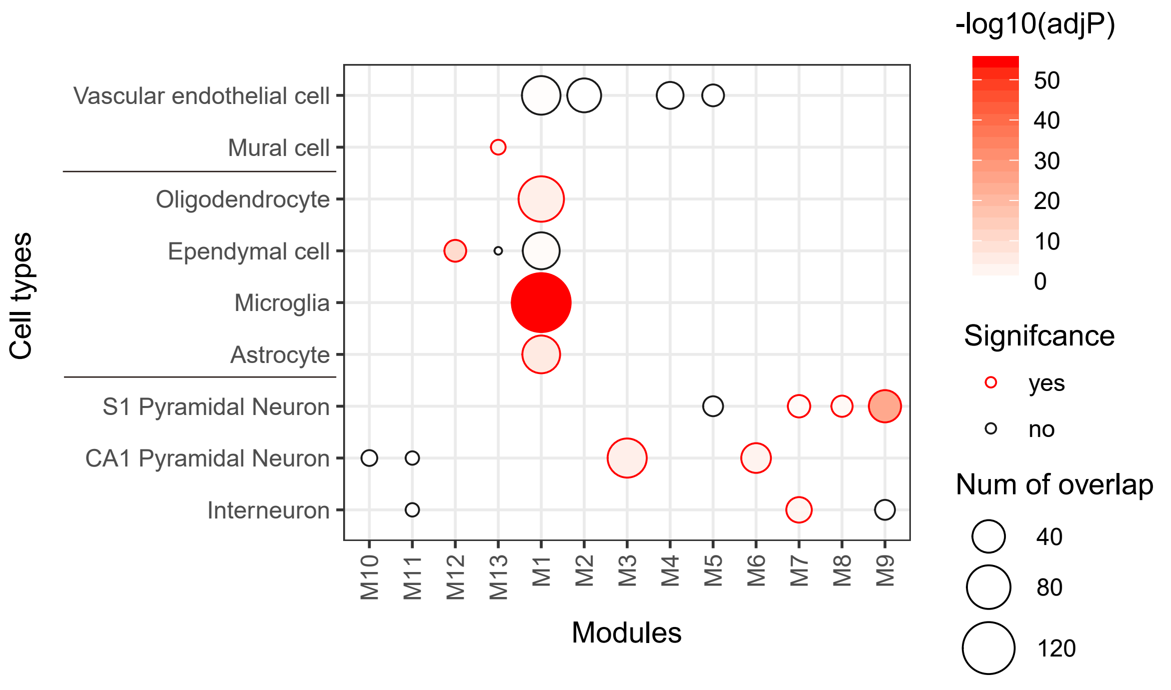

### Supplementary Fig. S4

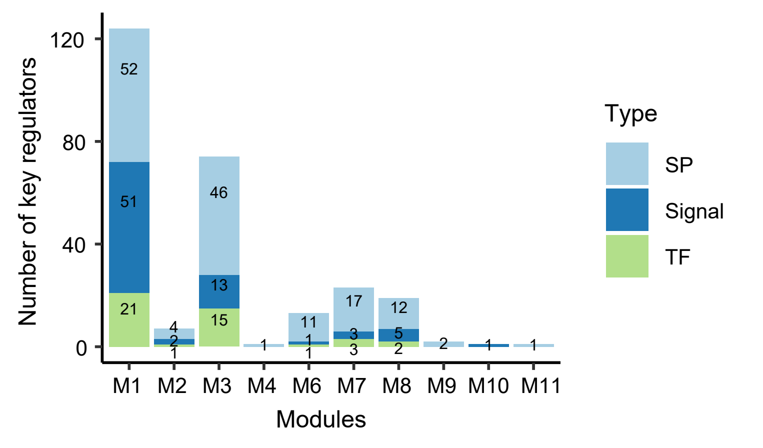

### Supplementary Fig. S5

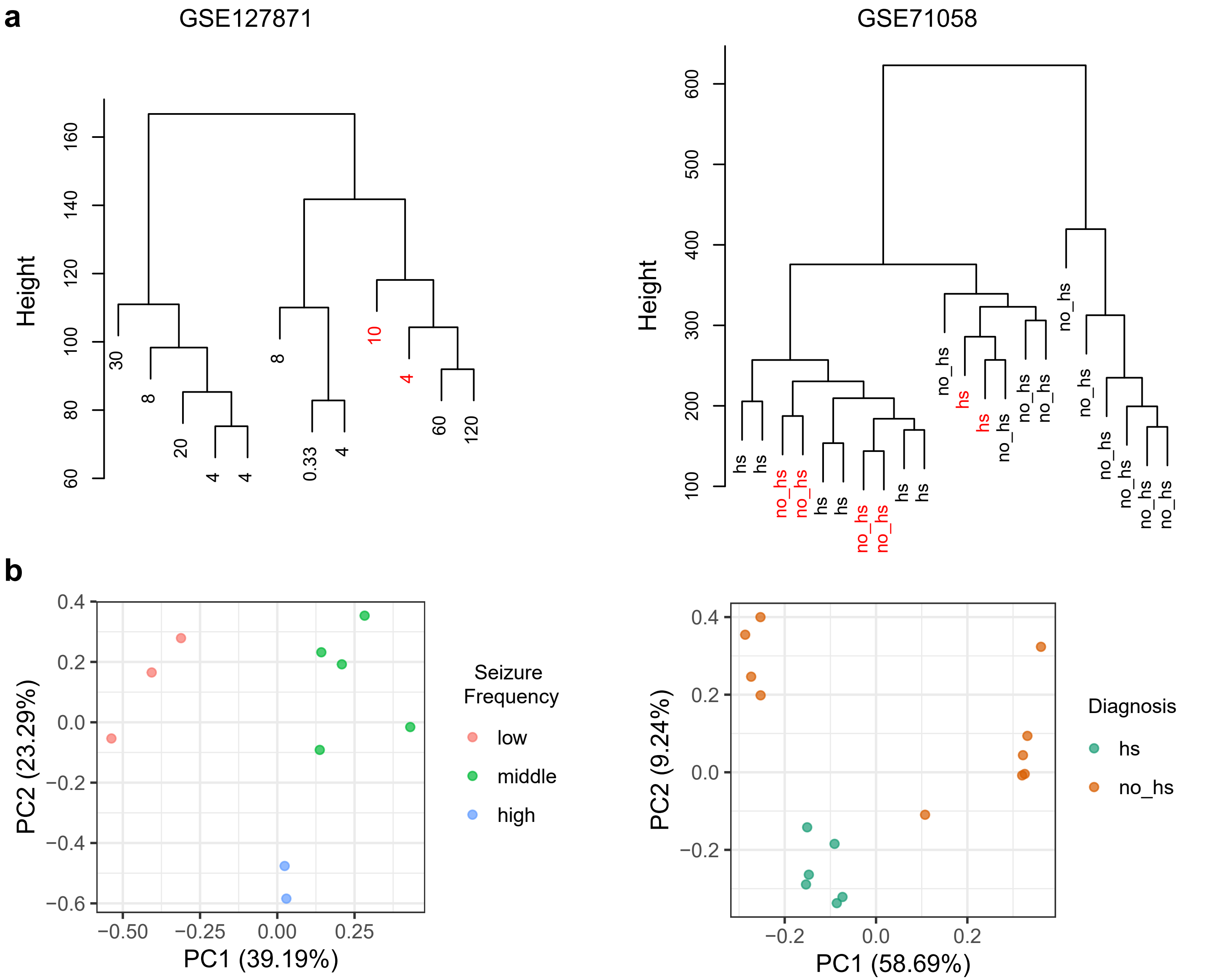

### Supplementary Fig. S6

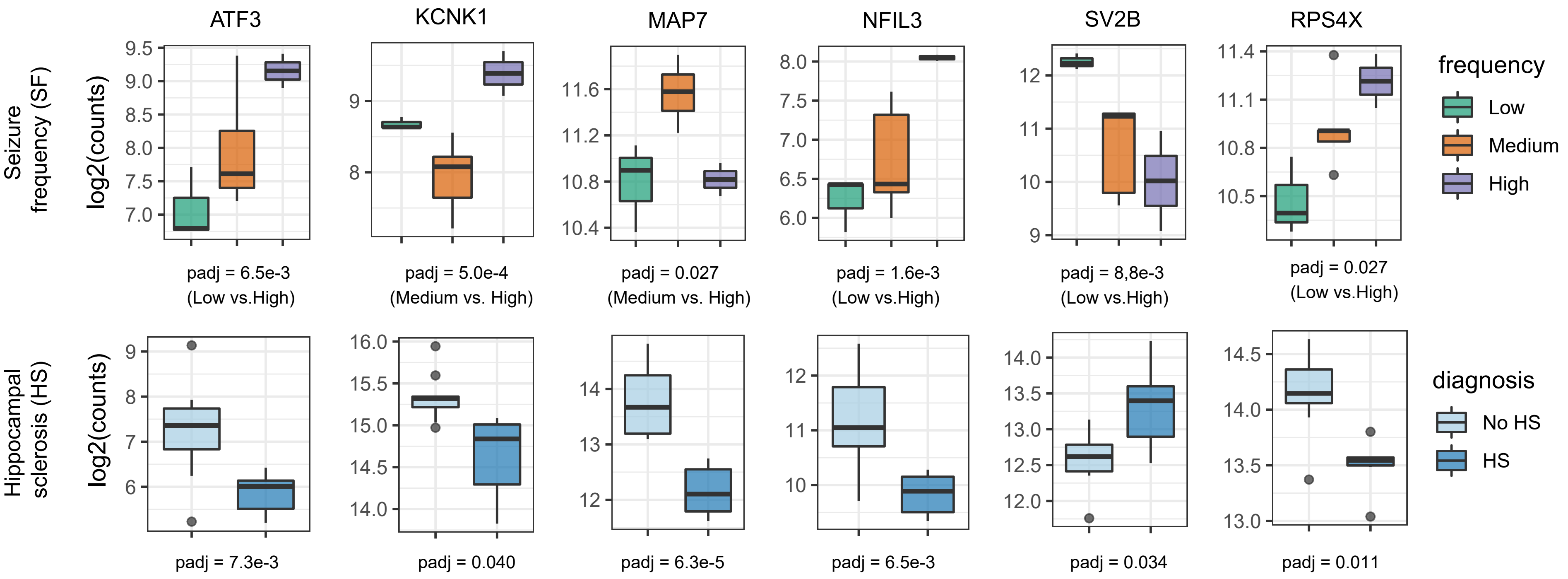
